# A Clinically Silent Resistance Phenotype That Promotes *Acinetobacter baumannii* Survival During Colistin Therapy

**DOI:** 10.64898/2026.01.20.700680

**Authors:** Muneer Yaqub, Namrata Bonde, Tuhina Maity, Ravali Arugonda, Zoe Du, Chinmai Rudra, Suman Tiwari, Roberto Jhonatan Olea-Ozuna, Sinjini Nandy, Joseph Boll, Jonathan Monk, Nicholas Dillon

## Abstract

*Acinetobacter baumannii* is a major cause of multidrug-resistant nosocomial infections, particularly ventilator-associated pneumonia, for which therapeutic options are increasingly limited. Colistin, a polymyxin antibiotic, is a drug of last resort for *A. baumannii,* boasting high susceptibility rates. Yet, despite relatively low rates of breakpoint-defined colistin resistance, clinical outcomes are highly variable, and the bacterial strategies that enable survival during colistin therapy remain poorly understood. Here, we integrate supervised machine-learning–guided genomic prioritization with functional, physiological, and in vivo analyses to interrogate the genetic basis of colistin response in *A. baumannii*. Machine-learning analysis of clinical isolates identified candidate loci associated with colistin survival, many of which did not alter minimum inhibitory concentration (MIC) when disrupted. Instead, growth-dynamic assays uncovered a subset of mutants capable of maintaining fitness upon inhibitory colistin exposure despite classification as susceptible via standardized antibiotic susceptibility testing. We define this phenotype as clinically silent resistance (CSR), a genetically encoded, MIC-independent survival state. Using a murine pneumonia model, we further demonstrate that CSR mutants thrive during colistin therapy in vivo. Together, these findings reveal a hidden layer of colistin survival that is not captured by standard susceptibility testing and highlight fundamental limitations of breakpoint-centric paradigms for predicting treatment outcomes in *A. baumannii*.

## Introduction

The global rise of multidrug-resistant *Acinetobacter baumannii* has placed increasing pressure on last-line antibiotics, with colistin frequently serving as a final therapeutic option for severe hospital-acquired infections, including ventilator-associated pneumonia and bloodstream infections [1–3]. Despite its clinical importance, outcomes of colistin therapy remain highly variable, and treatment failure is frequently observed even when isolates are classified as susceptible by standard antimicrobial susceptibility testing [4,5]. Understanding the bacterial strategies that enable survival under colistin pressure therefore remains a critical challenge for both antimicrobial resistance biology and clinical practice [6,7].

Colistin exerts bactericidal activity primarily through electrostatic interactions with lipopolysaccharide in the outer membrane, leading to membrane destabilization, depolarization, and cell death [8]. Canonical resistance mechanisms – including lipid A modification or complete lipopolysaccharide loss – directly elevate minimum inhibitory concentrations (MICs) as measured by Antimicrobial Susceptibility Testing (AST) and form the basis of current diagnostic frameworks [9–11]. While these mechanisms are well characterized, they do not fully explain the heterogeneity of clinical outcomes, nor do they account for infections that persist or recur despite apparent susceptibility [12,13]. These discrepancies suggest that bacterial survival under colistin exposure may extend beyond breakpoint-defined resistance alone [14].

Across bacterial pathogens, diverse non-canonical survival strategies – such as tolerance, persistence, and heteroresistance – have highlighted the limitations of static susceptibility testing in predicting therapeutic success [15–17]. However, these phenomena are often studied independently and remain incompletely integrated into clinical paradigms. In *A. baumannii*, the extent to which genetically encoded survival states operate independently of canonical resistance mechanisms – and whether such states meaningfully influence infection outcomes during colistin therapy – remains poorly defined.

Large-scale bacterial genomics offers a powerful avenue to interrogate resistance-associated features beyond well-defined mechanisms [18]. However, conventional genome-wide association approaches face substantial challenges in the context of colistin resistance. Resistant isolates are relatively rare, adaptive traits may be polygenic and heterogeneous, and survival-associated features are frequently embedded within conserved genes rather than acquired through horizontal gene transfer [19–21]. Under these conditions, classical statistical frameworks are underpowered to detect low-frequency, context-dependent signals that may nonetheless be biologically meaningful.

In parallel, machine-learning approaches have increasingly been adopted across the biological sciences to address complex questions characterized by high-dimensional variation, weak individual effects, and non-linear interactions [22–24]. Rather than testing single loci in isolation, supervised learning frameworks enable prioritization of genomic features associated with phenotypic outcomes across entire datasets, even when traditional assumptions of independence and effect size are violated. In microbial genomics, such approaches are now widely used to complement classical inference by generating experimentally testable hypotheses about resistance, virulence, and adaptation – particularly in settings where canonical approaches fall short [25,26].

Here, we applied a supervised machine-learning framework to population-scale *A. baumannii* genomic data to prioritize genetic features associated with clinical colistin resistance, and combined this with systematic functional genetics, physiological profiling, and in vivo infection modeling to assess the influence of the predications on responses to colistin. Through this integrative approach, we identify clinically silent resistance (CSR) – a genetically encoded, MIC-independent survival state that manifests during colistin therapy despite apparent susceptibility.

Unlike other non-canonical survival phenotypes such as tolerance, persistence, or heteroresistance – which are typically transient, stochastic, or confined to rare subpopulations – CSR represents a stable, genetically defined survival program that is expressed under host-relevant conditions without altering standard susceptibility measurements. Consequently, CSR remains invisible to breakpoint-based diagnostics yet contributes meaningfully to treatment failure in vivo. This work expands current paradigms of antimicrobial resistance and highlights fundamental limitations of breakpoint-centric diagnostics in predicting treatment outcome.

## Results

### Machine-learning–guided identification of candidate loci associated with colistin response

To identify genetic determinants associated with colistin resistance, we curated 310 *A. baumannii* clinical genomes from public repositories together with corresponding colistin susceptibility classifications from clinical testing. Because isolates classified as colistin resistant under CLSI guidelines are relatively rare in available datasets, conventional genome-wide association approaches are inherently underpowered to detect low-frequency but functionally important resistance-associated features [27,28] (**Figure 1a**). We therefore employed a supervised machine-learning framework to identify genomic patterns associated with colistin resistance across the full dataset, enabling systematic prioritization of candidate loci for downstream experimental validation.

**Figure 1.**
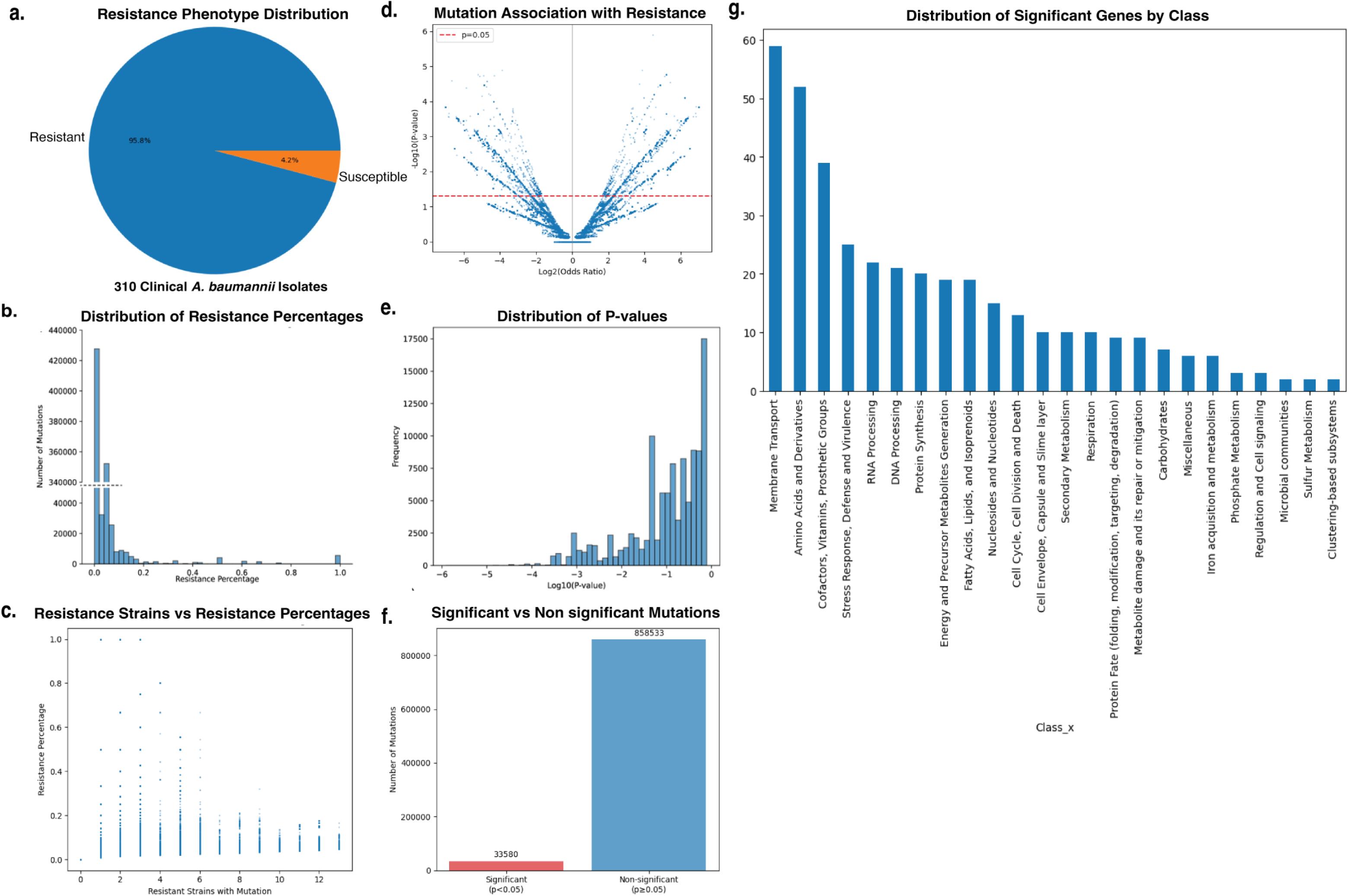
Machine-learning–guided identification of resistance-associated loci in *A. baumannii*. **(a)** Distribution of colistin susceptibility phenotypes across the curated clinical isolate dataset, illustrating the pronounced class imbalance between susceptible and resistant strains that limits the power of frequency-based association approaches. **(b)** Distribution of resistance percentages across all detected mutations. A histogram showing the frequency distribution of resistance percentages among the analyzed mutations, with a break in the y-axis (number of mutations) between 40,000 and 430,000 to accommodate the large number of mutations that occur infrequently in resistant isolates. **(c)** Relationship between mutation recurrence and resistance specificity. A scatter plot showing the relationship between the number of resistant strains carrying a given mutation and the corresponding resistance percentage, demonstrating that mutation recurrence among resistant isolates does not necessarily confer resistance specificity. **(d)** Association strength of individual mutations with colistin resistance. A volcano plot illustrating the relationship between mutation effect size (log₂ odds ratio) and statistical significance (−log₁₀ *P* value). The red dashed line indicates the nominal significance threshold ( *P* = 0.05), highlighting the sparse subset of mutations exhibiting preferential association with resistance amid extensive background variation. **(e)** Global distribution of statistical support for mutation–resistance associations. A histogram displaying the distribution of *P* values (<1) on a logarithmic scale (log₁₀ *P* value), showing enrichment of low *P* values relative to uniform expectation while indicating that the majority of mutations remain non-significant. **(f)** Summary of statistically significant versus non-significant mutation associations. A bar chart comparing the total number of mutations classified as significant (*P* < 0.05) and non-significant (*P* ≥ 0.05), with counts annotated above each bar. **(g)** Functional classification of genes harboring statistically prioritized mutation associations, revealing enrichment across diverse cellular processes rather than exclusive dominance of canonical resistance pathways.

We reasoned that resistance-associated genes would most plausibly be distinguished by patterns of allelic variation. Accordingly, the machine-learning framework first evaluated the global distribution of mutations across the dataset to identify loci enriched for variation associated with resistant phenotypes. This analysis revealed that the vast majority of genes carried few or no resistance-associated mutations, whereas a smaller subset harbored multiple mutations across isolates (**Figure 1b**). However, mutation enrichment alone proved insufficient for prioritization: many mutations occurred predominantly in susceptible backgrounds, and even mutations observed in multiple resistant isolates rarely showed strong resistance specificity (**Figure 1c**).

We therefore evaluated whether resistance-associated mutations could instead be distinguished by the strength of their association with resistant phenotypes. While a small subset of mutations exhibited elevated effect sizes and statistical support (**Figure 1d**), the global distribution of p-values revealed that such signals were rare relative to the full mutational landscape (**Figure 1e**), with only a small fraction of mutations meeting significance thresholds (**Figure 1f**).

Next, we assessed whether resistance-associated signals could instead be explained by differential gene content. Gene presence–absence analysis revealed that, with few exceptions, candidate loci were broadly shared between resistant and susceptible isolates. Only a small number of genes exhibited enrichment or depletion in resistant genomes **(Supplementary Table 1)**,

Because neither mutation distribution nor gene presence–absence alone was sufficient to explain resistance-associated patterns, we incorporated functional context as an additional criterion for gene prioritization. Functional annotation of loci implicated by the machine-learning framework revealed enrichment across diverse cellular processes, including membrane transport, metabolism, stress-response pathways, and regulatory functions (**Figure 1g**). This functional heterogeneity suggested that colistin resistance reflects distributed physiological adaptations rather than isolated canonical mechanisms.

On this basis, we adopted an intentionally inclusive gene selection strategy for experimental validation. Rather than prioritizing loci solely by statistical rank or mutation burden, we integrated mutation-level signals with functional classification to sample broadly across resistance-associated biology. Candidate genes were selected to include loci both with and without enriched mutation signatures, and to ensure representation across multiple functional subsystems. Using this functionally stratified framework, we prioritized 29 genes spanning eleven cellular subsystems for downstream experimental interrogation (**Table 1; Extended Table 1)**.

**Table 1:**
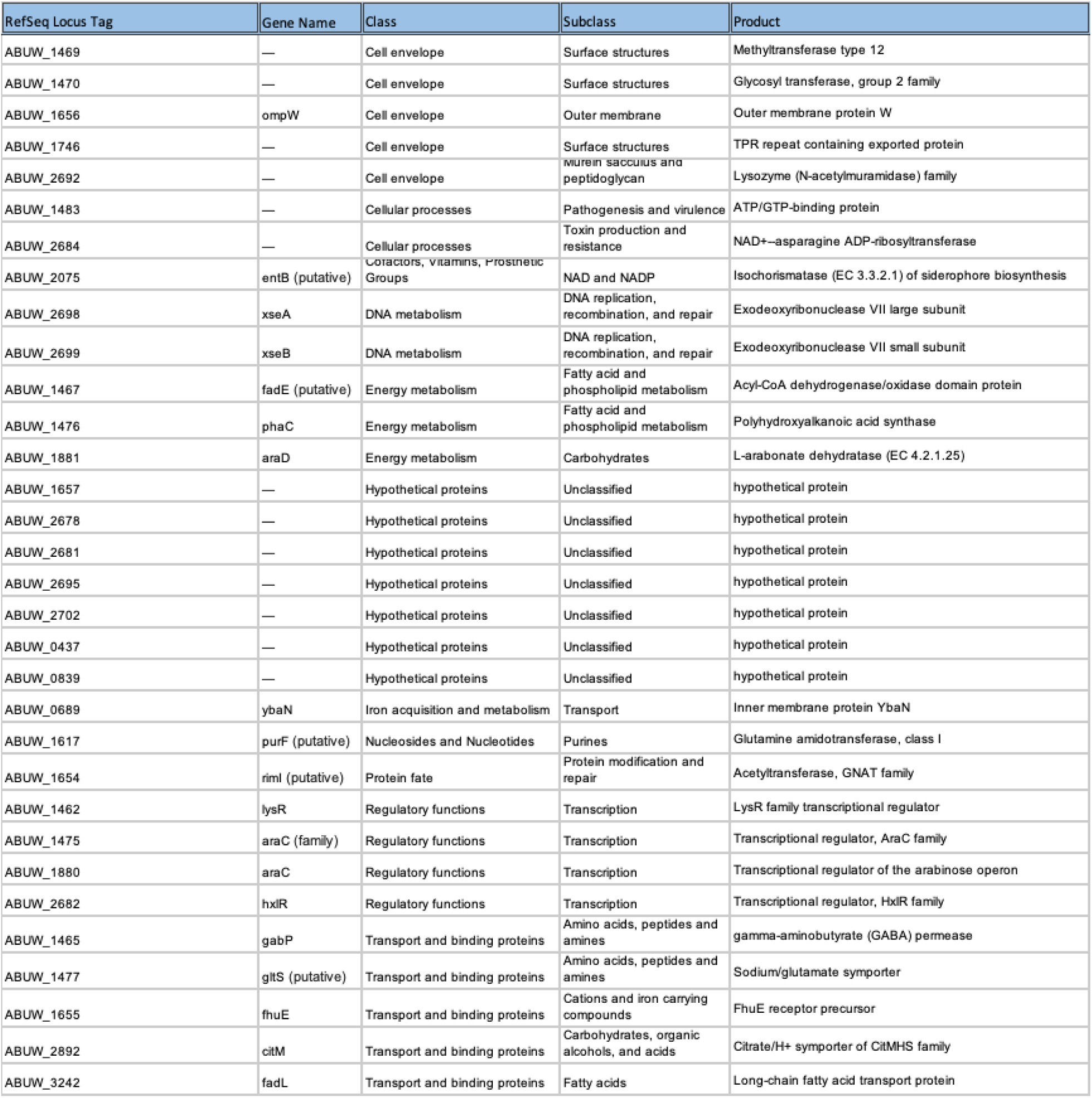
Functional Analysis of Machine-learning–prioritized genes. Table summarizes the *A. baumannii* loci prioritized by supervised machine learning and selected for experimental interrogation using transposon mutants in the AB5075 background. For each gene, the table lists the locus identifier, predicted or annotated function, mutation-level association statistics derived from population-genomic analysis, and the number of resistant clinical isolates harboring mutations at that locus.

**Table 2:** Summary of growth-kinetic phenotypes across machine-learning–prioritized mutants. Classification of mutants based on growth behavior under colistin stress, including wild-type–like inhibition, partial growth recovery, and Clinically Silent Resistant phenotypes.

| Invariant | Partial | Clinically Silent Resistant |
| --- | --- | --- |
| ABUW_1470::Tn | ABUW_3242::Tn | ABUW_1465::Tn |
| ABUW_2698::Tn | ABUW_1617::Tn | ABUW_2892::Tn |
| ABUW_1475::Tn | ABUW_2681::Tn | ABUW_1483::Tn |
| ABUW_1655::Tn | ABUW_1462::Tn |  |
| ABUW_1476::Tn | ABUW_1477::Tn |  |
| ABUW_2684::Tn | ABUW_1657::Tn |  |
| ABUW_2682::Tn | ABUW_2702::Tn |  |
| ABUW_1880::Tn | ABUW_1469::Tn |  |
| ABUW_0689::Tn | ABUW_2075::Tn |  |
| ABUW_1746::Tn | ABUW_1881::Tn |  |
| ABUW_1467::Tn |  |  |
| ABUW_1656::Tn |  |  |
| ABUW_2678::Tn |  |  |
| ABUW_1654::Tn |  |  |
| ABUW_2692::Tn |  |  |
| ABUW_0839::Tn |  |  |

### MIC testing does not detect altered colistin susceptibility in machine-learning–prioritized *A. baumannii* mutants

To assess whether machine-learning–prioritized genes directly alter categorical colistin susceptibility, we employed a transposon mutant library constructed in *A. baumannii* AB5075-UW (ABUW), a genetically tractable reference clinical strain that is phenotypically susceptible to colistin yet retains intact physiological stress-response and adaptive survival pathways. We obtained corresponding transposon insertion mutants from the Manoil library and validated insertion sites by PCR [29] **(Extended Figure 1)**. Each mutant was subjected to standardized broth microdilution AST to determine the minimum inhibitory concentration (MIC) of colistin relative to the wild-type strain. Across the mutant panel, none exhibited MIC values distinguishable from wild type (**Figure 2**).

**Figure 2:**
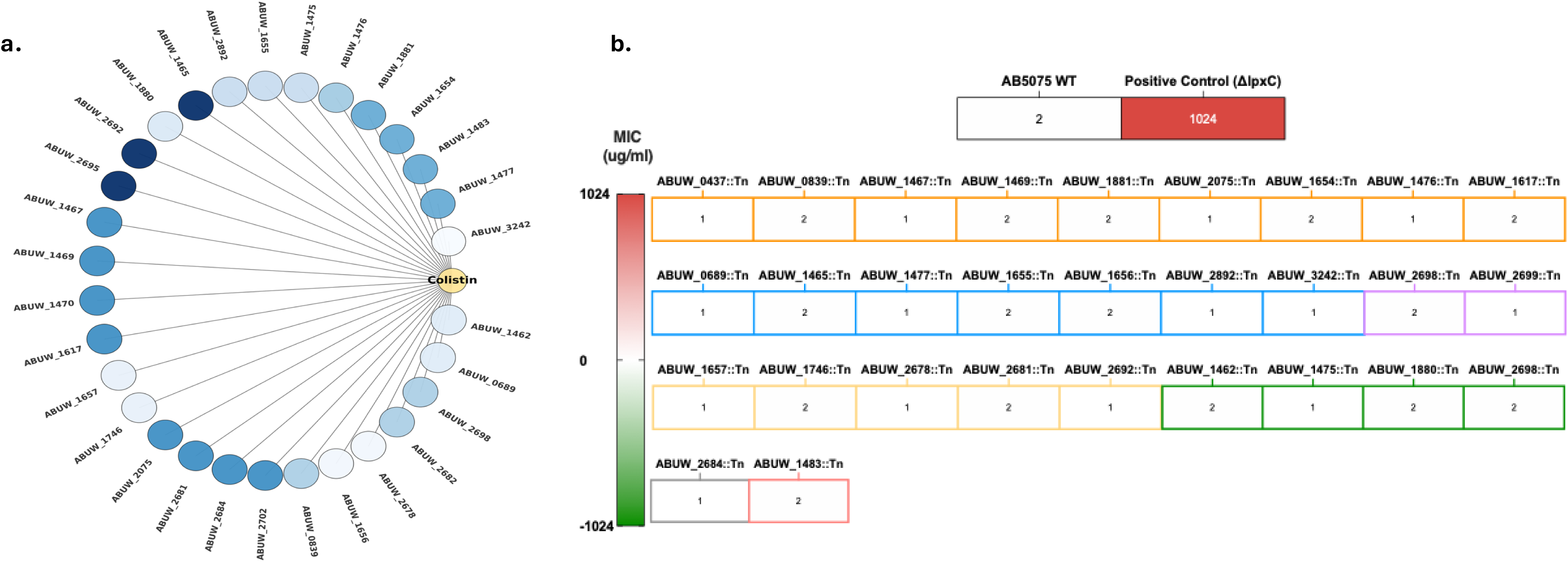
MIC testing does not detect altered colistin susceptibility in machine-learning–prioritized mutants. Minimum inhibitory concentrations (MICs) of colistin for transposon mutants in *A. baumannii* AB5075 compared with wild type, determined by broth microdilution. All tested mutants exhibited MIC values indistinguishable from wild type, indicating no shift in categorical susceptibility.

Thus, despite population-genomic associations with colistin response observed in clinical isolates, loss-of-function disruption of these loci did not produce shifts in categorical susceptibility based on established clinical breakpoints. These results indicate that standard MIC testing did not detect changes in categorical colistin susceptibility for the machine-learning–prioritized mutants.

### Growth dynamics reveal a clinically silent resistance phenotype

Although MIC testing did not reveal changes in categorical colistin susceptibility, we reasoned that disruption of machine-learning–prioritized loci might influence bacterial survival in ways not captured by binary breakpoint-based assays [30]. To test this possibility, we quantified real-time growth dynamics for each mutant under increasing colistin concentrations, enabling detection of adaptive fitness differences beyond static MIC thresholds.

When assessed across a gradient of 0–4 µg ml⁻¹ colistin, the wild-type strain exhibited complete growth inhibition at 2 µg ml⁻¹, consistent with its MIC value (**Figure 3a, Extended Figure 3)**. Approximately half of the mutants (n = 16) displayed similar kinetic suppression, indicating no measurable fitness advantage under antibiotic stress (**Figure 3b**). In contrast, distinct growth behaviors emerged among the remaining mutants that were not explained by their MIC profiles.

**Figure 3:**
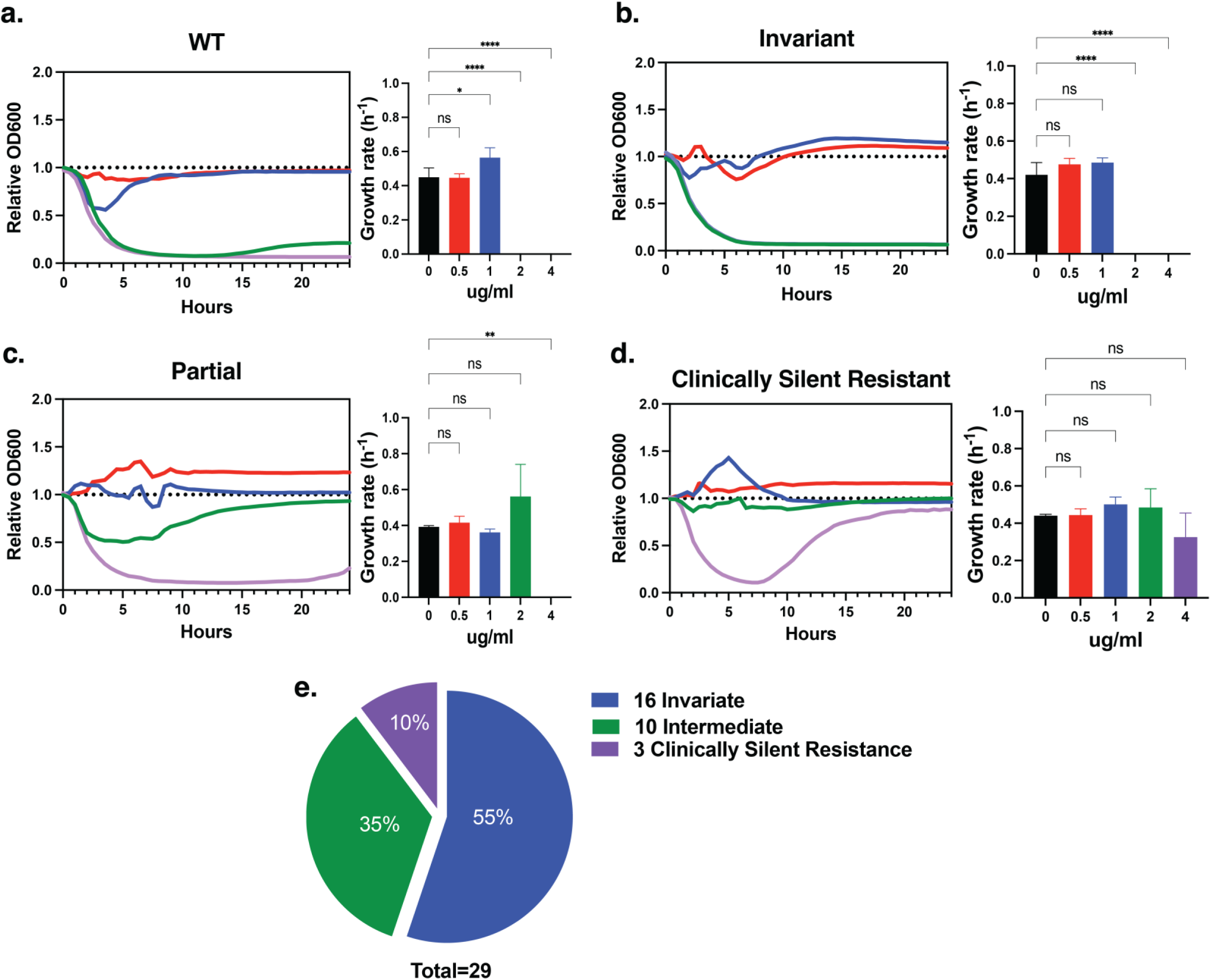
Growth dynamics reveal a clinically silent resistance phenotype. (a) Growth kinetics of wild-type *A. baumannii* AB5075 across increasing colistin concentrations (0–4 µg ml⁻¹). (b) Representative growth curves of mutants exhibiting wild-type–like kinetic suppression under colistin exposure. (c) Growth kinetics of intermediate mutants displaying partial growth recovery at 2 µg ml⁻¹ colistin, a concentration inhibitory to wild type. (d) Growth kinetics of clinically silent resistant (CSR) mutants exhibiting sustained or recovered exponential-phase growth at inhibitory colistin concentrations, including continued growth at 4 µg ml⁻¹. (e) Pie chart summarizing the distribution of growth phenotypes across all 29 tested mutants, categorized as wild-type–like, intermediate (2 µg ml⁻¹), or clinically silent resistant.

Ten mutants maintained partial growth at 2 µg ml⁻¹ colistin, suggesting modest adaptive capacity at concentrations inhibitory to wild type, whereas three mutants continued to actively replicate even at 4 µg ml⁻¹ colistin, as evidenced by sustained increases in optical density during prolonged incubation (**Figure 3c–e**). Importantly, although these cultures exhibited measurable growth, total biomass accumulation remained well below the ≥90% inhibition threshold used to define resistance in standardized minimum inhibitory concentration (MIC) assays. As a result, these mutants remained classified as colistin susceptible by conventional antimicrobial susceptibility testing despite exhibiting replication under drug pressure.

These three mutants – disrupting *ABUW_2892 (citN),* encoding a citrate transporter; *ABUW_1483*, a putative AAA⁺ ATPase; and *ABUW_1465 (gabP),* a γ-aminobutyrate permease – were designated as the clinically silent resistant (CSR) group (**Table 1; Extended Table 2)**. Despite their diverse functional annotations, all three shared a defining phenotype: the capacity of a subpopulation to recover and sustain replication during prolonged colistin exposure without achieving the population-level growth required to exceed MIC-defined resistance thresholds. This behavior represents a genetically encoded, MIC-independent survival state that is not captured by categorical susceptibility paradigms.

This distinction reflects fundamental differences in how susceptibility is operationally defined. Standard MIC determinations assess whether antibiotic exposure suppresses population-level growth beyond a fixed categorical threshold, whereas growth-curve analyses resolve whether any fraction of the population retains the capacity to replicate under sustained antibiotic stress. The CSR phenotype emerges within this diagnostic gap, in which low-level resistance is detectable by kinetic growth measurements yet remains invisible to MIC-based classification.

Notably, CSR-associated genes spanned a broad range of mutation-level statistical support in population-genomic analyses **(Extended Table 2)**. Whereas *gabP (ABUW_1465)* exhibited borderline enrichment for resistance-associated mutations, *citN (ABUW_2892)* and *ABUW_1483* showed no significant mutation-level association. Thus, the emergence of CSR phenotypes could not be reliably predicted from population-scale genomic statistics alone, underscoring the need for targeted physiological profiling to identify adaptive survival states that escape conventional susceptibility diagnostics.

### Physiological profiling reveals widespread stress adaptation distinct from clinically silent resistance

We next performed a stepwise physiological profiling of 29 AB5075 transposon mutants under colistin-relevant conditions to determine whether machine-learning–prioritized loci influence *A. baumannii* physiology in ways not captured by MIC testing. Because colistin primarily disrupts the bacterial outer membrane [31], we first examined envelope integrity using a dual-dye permeability assay. Membrane permeability was assessed with propidium iodide, which penetrates only cells with compromised membranes and fluoresces upon binding nucleic acids, while membrane polarization was measured using 3,3′-diethyloxacarbocyanine iodide (DiOC₂(3)), a voltage-sensitive dye that accumulates in cells in a membrane potential–dependent manner and exhibits a fluorescence shift upon depolarization [32].

Only a small subset of mutants exhibited increased membrane permeability relative to wild type (2/29), accompanied by significant membrane depolarization in the same limited number of strains (**Figure 4a,b**). These defects were observed in mutants disrupting *ABUW_1656 (ompW)* and *ABUW_1475 (araC),* both of which fall within the Invariant category, as well as *ABUW_2075* (2,3-dihydro-2,3-dihydroxybenzoate synthetase), which exhibited a Partial growth phenotype. Together, these findings indicate that both envelope-associated components and upstream metabolic or regulatory factors contribute to membrane stability during colistin exposure.

**Figure 4:**
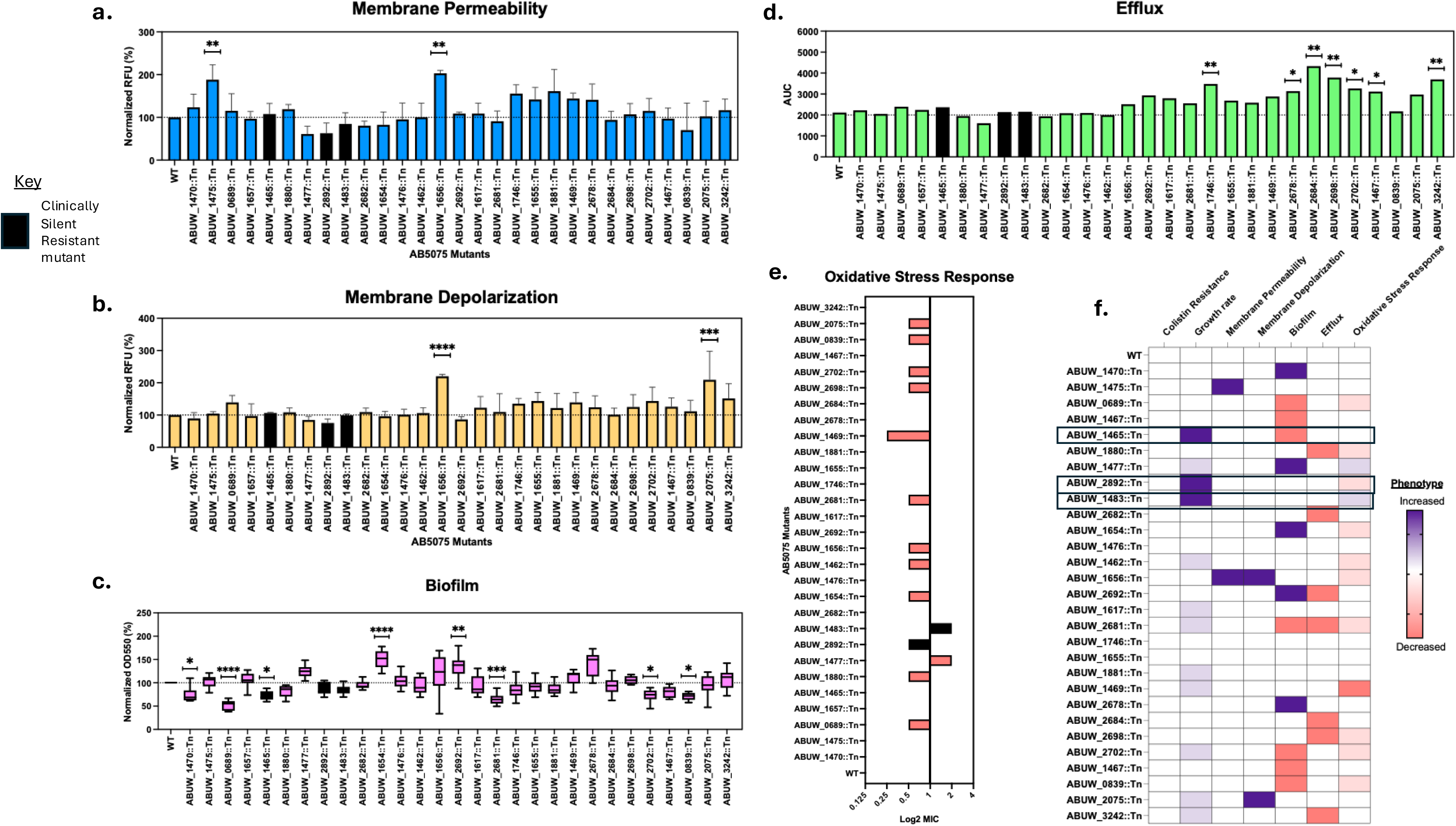
Physiological profiling reveals stress adaptation distinct from clinically silent resistance. Physiological responses of machine-learning–prioritized *A. baumannii* mutants were quantified under colistin-relevant conditions and compared with wild type. (a) Outer-membrane permeability, measured as relative uptake of a dual-dye fluorescent probe and expressed as fold change relative to wild type. (b) Membrane potential, quantified using a depolarization-sensitive fluorescent dye and reported as normalized fluorescence intensity. (c) Efflux capacity, assessed by ethidium bromide accumulation and plotted as relative intracellular fluorescence, where higher values indicate reduced efflux activity. (d) Biofilm formation, measured by crystal-violet staining and expressed as absorbance normalized to wild type. (e) Oxidative-stress tolerance, quantified as relative survival following hydrogen peroxide challenge. (f) Summary heatmap showing deviations of each mutant from wild-type behavior across all physiological assays, with values scaled and centered per assay. Data represent means ± s.d. from at least three independent biological replicates. Statistical significance was determined by comparison to wild type using t-test.

Because biofilm formation is a known modulator of colistin tolerance [33,34], we quantified biofilm production using a crystal-violet assay. Biofilm capacity was altered in 12/29 mutants, with 5/29 displaying increased biomass and 7/29 exhibiting reduced biofilm formation relative to wild type (**Figure 4c**). These effects mapped to genes involved in metabolism, envelope modification, and transport, indicating that machine-learning–prioritized loci broadly reshape biofilm-associated physiology during antibiotic exposure. Among these, a single CSR mutant, ABUW_1465::Tn (*gabP*), exhibited increased biofilm formation, whereas the remaining biofilm-altered mutants were distributed across the Partial (3/12) and Invariant (8/12) growth categories.

We next assessed intracellular antibiotic handling by quantifying efflux capacity using ethidium bromide accumulation assays. A subset of mutants (7/29) exhibited impaired efflux, reflected by increased intracellular fluorescence (**Figure 4d**). These mutants spanned multiple functional categories, including metabolic enzymes, regulatory proteins, and hypothetical proteins, indicating that efflux modulation during colistin stress arises from convergence across diverse cellular pathways rather than disruption of a single efflux system. Notably, none of the efflux-impaired mutants belonged to the clinically silent resistant (CSR) group; instead, they were distributed across the Invariant (5/7) and Partial (2/7) growth categories.

Finally, given evidence that colistin induces oxidative stress [35], we examined redox tolerance using hydrogen peroxide challenge assays. Oxidative stress responses were perturbed in 14/29 mutants, displaying a pronounced asymmetry: only 2/29 mutants exhibited increased tolerance, whereas 12/29 were sensitized (**Figure 4e**). These redox-altered mutants were evenly distributed between the Partial (6/14) and Invariant (6/14) growth categories. Notably, two CSR mutants exhibited altered redox phenotypes, with ABUW_1483::Tn displaying increased oxidative stress tolerance and ABUW_2892::Tn (*citN*) exhibiting sensitization, indicating that modulation of oxidative stress handling is a shared axis engaged by a subset of clinically silent resistant strains.

Collectively, most machine-learning–prioritized mutants (24/29) deviated from wild-type behavior in at least one physiological dimension (**Figure 4f**). Biofilm formation and oxidative stress tolerance were the most frequently perturbed phenotypes, whereas direct membrane destabilization and energetic collapse were comparatively rare. Notably, the three CSR mutants – disrupting *ABUW_2892 (citN), ABUW_1483*, and *ABUW_1465 (gabP)* – converged on a limited set of physiological axes but through distinct routes: two engaged oxidative stress responses with opposing outcomes, while the third adopted a biofilm-associated survival state. These data indicate that clinically silent resistance is supported by alternative network configurations involving redox and biofilm-associated physiology rather than a single dominant resistance mechanism.

Consistent with this axis-specific engagement, the functional identities of CSR loci further distinguish them from invariant and partial mutants. Whereas invariant and partial mutants reflected widespread perturbation of structural and regulatory functions – distributed across cell envelope components, transcriptional regulators, metabolic enzymes, and hypothetical proteins – the CSR loci were biased toward transport and intracellular homeostasis. *ABUW_2892 (citN)* and *ABUW_1465 (gabP)* encode inner-membrane transporters involved in organic acid and amino acid flux, respectively, while *ABUW_1483* encodes a putative ATP/GTP-binding protein associated with cellular processes and virulence-related functions. Notably, none of the CSR genes encode outer-membrane components, lipid A biosynthesis enzymes, or canonical regulatory factors typically implicated in colistin resistance.

Moreover, this functional divergence mirrored population-genomic signals from our machine learning analysis: the biofilm-associated CSR mutant (*gabP*) harbored unique resistance-associated mutations, whereas the redox-engaged CSR mutants did not **(Extended Table 1)**. Together, these observations suggest that clinically silent resistance preferentially emerges through metabolic and transport-linked rewiring that modulates intracellular stress handling—particularly redox balance and biofilm-associated states – without invoking classical envelope remodeling or breakpoint-defined resistance mechanisms.

### In vivo pneumonia model reveals clinically silent resistance under colistin therapy

We next employed a murine pneumonia model of *A. baumannii* infection, in which bacteria were delivered intratracheally into the lungs [36], to determine if the CSR mutants impacted colistin treatment outcomes in a host (**Figure 5a**). We first performed a dose–response analysis across 0–20 mg kg⁻¹, administered intraperitoneally one hour after infection with the AB5075 wild-type strain, to establish the therapeutic window of colistin in this model. Colistin produced a clear dose-dependent reduction in pulmonary bacterial burden, with 5 mg kg⁻¹ resulting in an approximately 1–1.5 log₁₀ reduction in lung CFU (from ∼10⁷ to ∼5 × 10⁵ CFU) and 10 mg kg⁻¹ achieving an approximately 4 log₁₀ reduction (to ∼10³ CFU) relative to untreated controls, representing the lowest doses that effectively suppressed infection (**Figure 5b**).

**Figure 5:**
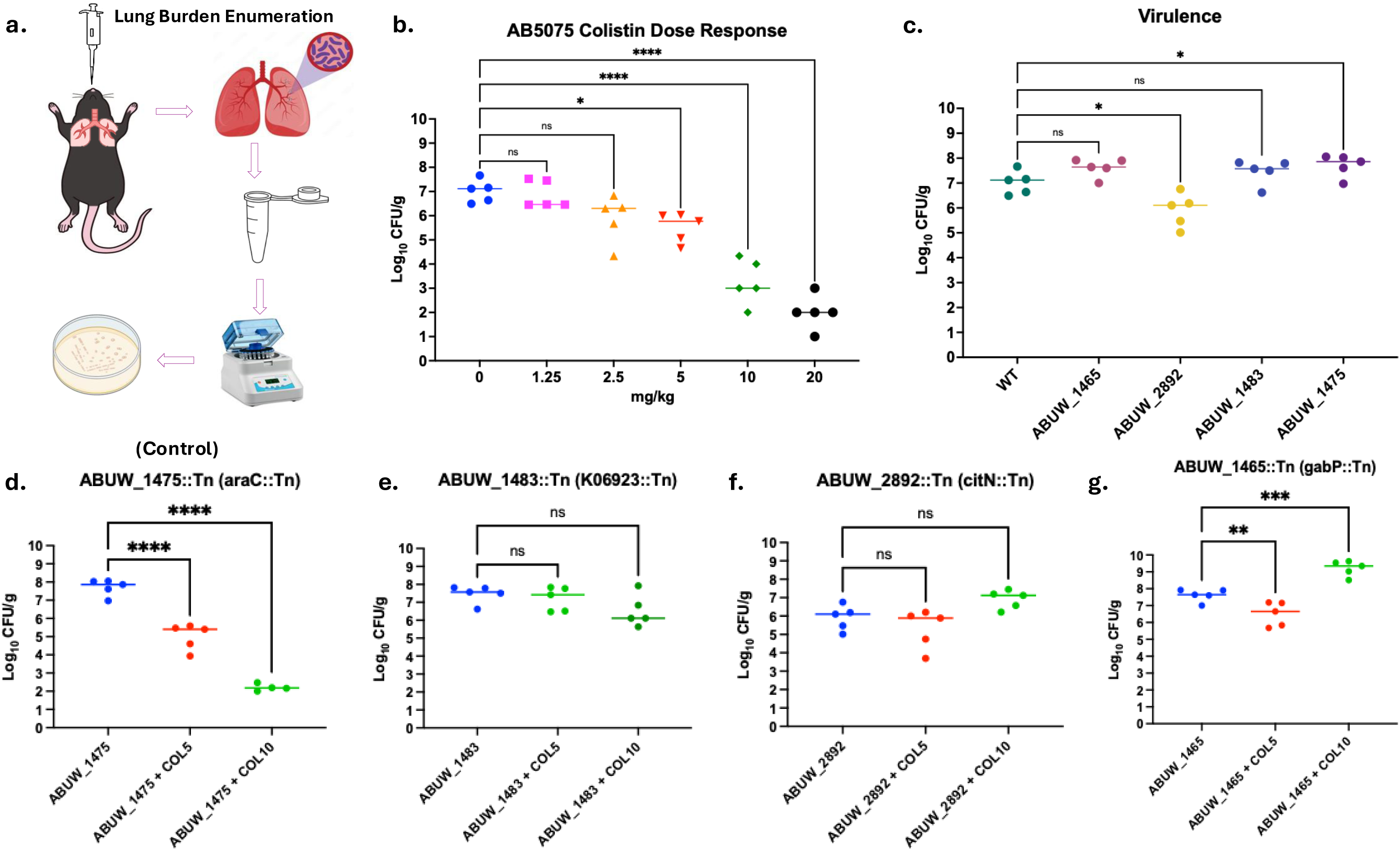
In vivo pneumonia model reveals clinically silent resistance under colistin therapy. In vivo survival of *A. baumannii* strains was assessed using a murine pneumonia model under colistin treatment. (a) Schematic overview of the experimental design, including intratracheal infection, timing of colistin administration, and bacterial burden assessment. (b) Dose–response of colistin efficacy against wild-type *A. baumannii* AB5075 in the lung, plotted as colony-forming units (CFU) per lung following treatment with increasing colistin doses. (c) Pulmonary bacterial burdens (CFU per lung) for wild type, a non-advantaged control mutant, and clinically silent resistant (CSR) mutants in untreated mice, indicating baseline infection outcomes. (d) Lung bacterial burdens following colistin treatment of mice infected with wild type or the non-advantaged control mutant. (e–g) Lung bacterial burdens following colistin treatment of mice infected with individual CSR mutants, showing persistence or expansion despite therapy. Each data point represents an individual animal; horizontal lines denote group medians. Statistical comparisons were performed relative to wild type using non-parametric tests appropriate for CFU distributions, with multiple-comparison correction where applicable. Sample sizes and exact *P* values are indicated in the figure.

We next compared infection outcomes for the wild type and an Invariant control mutant, ABUW_1475::Tn (*araC*), with those of the three mutants that exhibited CSR in vitro: ABUW_2892::Tn (*citN*), ABUW_1483::Tn, and ABUW_1465::Tn (*gabP*). In the absence of colistin, all strains established comparable pulmonary bacterial burdens, with lung CFU generally ranging from ∼10⁶ to 10⁸ across groups (**Figure 5c**). Differences between strains were modest, typically within ≤1 log₁₀, and did not follow a consistent directional pattern that distinguished CSR mutants from the invariant control or wild type. These data indicate that disruption of CSR-associated loci does not confer a baseline fitness advantage or defect during lung infection, suggesting that any differences observed during antibiotic therapy are unlikely to arise from enhanced virulence or growth in untreated hosts.

Next, we treated mice infected with each strain with either 5 mg kg⁻¹ or 10 mg kg⁻¹ colistin. In the invariant control mutant, ABUW_1475::Tn (*araC*), colistin treatment produced a robust, dose-dependent reduction in pulmonary bacterial burden, with an approximately 2.4 log₁₀ reduction at 5 mg kg⁻¹ and a ∼5.7 log₁₀ reduction at 10 mg kg⁻¹ relative to untreated infection, closely mirroring wild-type clearance.

In contrast, all three clinically silent resistant (CSR) mutants displayed attenuated responses to therapy that were inconsistent with their in vitro antimicrobial susceptibility profiles. ABUW_2892::Tn (*citN*) exhibited minimal suppression at both doses, with an approximately 0.2–0.3 log₁₀ reduction at 5 mg kg⁻¹ and no appreciable reduction at 10 mg kg⁻¹. ABUW_1483::Tn showed a modest, dose-dependent response, with little to no reduction at 5 mg kg⁻¹ and an approximately 1.5 log₁₀ reduction at 10 mg kg⁻¹, remaining substantially less responsive than wild type or the invariant control. Interestingly, ABUW_1465::Tn (*gabP*) exhibited a marked increase in pulmonary bacterial burden following treatment at 10 mg kg⁻¹, with lung CFU rising by approximately 1.7 log₁₀ relative to untreated infection (**Figure 5d–g**). Collectively, these data demonstrate that CSR mutants thrive in vivo under clinically relevant colistin dosing at bacterial burdens orders of magnitude higher than those observed for wild type or invariant controls.

These findings establish that clinically silent resistance extends beyond an in vitro artifact and constitutes a biologically meaningful survival strategy during infection. The capacity of CSR mutants to increase in burden during colistin therapy highlights a critical limitation of MIC-based susceptibility testing, revealing how genetically encoded yet diagnostically silent adaptations can compromise antibiotic efficacy and impede effective clearance of *A. baumannii* in vivo.

## Discussion

In this study, we integrate machine-learning–guided genomic prioritization with functional genetics, physiological profiling, and in vivo infection modeling to uncover a previously underappreciated mode of colistin survival in *A. baumannii*. While canonical resistance mechanisms elevate minimum inhibitory concentrations (MICs), and are readily detected by standard susceptibility testing, our findings demonstrate that a subset of genetically encoded adaptations enables bacterial survival under colistin exposure without altering categorical susceptibility. We define this phenomenon as clinically silent resistance (CSR) – a survival state that remains invisible to breakpoint-based diagnostics yet has tangible consequences for treatment efficacy in vivo.

Our machine-learning framework served as the origin point for this discovery by identifying candidate loci enriched for mutation patterns in colistin-resistant clinical isolates. These signals were frequently modest, heterogeneous, and distributed across conserved genes rather than driven by gene acquisition or loss, highlighting a polygenic and context-dependent architecture of colistin response. Importantly, the strength of population-genomic association did not reliably predict functional outcomes under experimental conditions. Loci with weak or absent mutation-level enrichment nevertheless gave rise to robust survival phenotypes, underscoring the limits of population-scale statistics for inferring antibiotic response in isolation.

Functional interrogation revealed a pronounced disconnect between MIC-defined susceptibility and bacterial survival. Disruption of most machine-learning–prioritized loci did not alter categorical susceptibility by MIC testing, confirming that these genes do not encode classical breakpoint-defining resistance determinants. However, growth-kinetics assays uncovered a subset of mutants capable of sustaining growth at colistin concentrations that fully inhibited the wild-type strain. These CSR mutants exhibited reproducible, MIC-independent survival advantages, demonstrating that growth dynamics provide a more sensitive and informative readout of antibiotic response than static MIC measurements alone.

Physiological profiling further clarifies how CSR occupies a diagnostic space that is invisible to conventional AST. Across the mutant panel, machine-learning–prioritized genes broadly perturbed multiple physiological axes – including efflux capacity, biofilm formation, and oxidative stress tolerance – yet these changes were heterogeneous and did not reliably translate into enhanced growth or survival under colistin exposure as defined by MIC-based criteria. In contrast, CSR mutants engaged a limited subset of physiological axes, most prominently oxidative stress handling or biofilm-associated states, but without producing the magnitude or coordination of phenotypic change required to elevate MICs. Thus, while canonical resistance mechanisms manifest as population-level growth sufficient to cross AST thresholds, CSR reflects survival-supportive network states that permit low level resistance below diagnostic cutoffs. This dissociation underscores a fundamental limitation of MIC-based frameworks: they are designed to detect overt resistance phenotypes but systematically overlook adaptive states that sustain bacterial survival during therapy.

The clinical relevance of CSR was most clearly demonstrated in a murine pneumonia model. Despite remaining classified as colistin-susceptible by MIC testing, CSR mutants thrived during colistin therapy in vivo, whereas wild-type bacteria and Invariant mutants were effectively suppressed. These findings provide a potential mechanistic explanation for a subset of unexplained treatment failures in colistin therapy and illustrate how diagnostically silent adaptations can undermine last-line antibiotic efficacy in host environments.

Several limitations of this study are important to consider, particularly with respect to the machine-learning framework that motivated downstream experimental analyses. First, model training was constrained by the relative scarcity of publicly deposited colistin-resistant clinical isolates, which limited statistical power at individual loci and reduced the ability to resolve higher-order interactions among mutations. As a result, many prioritized genes exhibited modest or heterogeneous mutation-level signals, reflecting the polygenic and context-dependent nature of colistin response rather than strong single-gene effects. Second, the models relied primarily on static genomic features, such as gene presence–absence and nucleotide substitutions, without incorporating regulatory, structural, or expression-level information. Survival states such as CSR may arise from network-level reprogramming that is not directly inferable from genome sequence alone, and the absence of transcriptomic or metabolomic features likely limited predictive resolution. Third, machine-learning outputs were intentionally treated as hypothesis-generating signals rather than definitive predictors of resistance. The emergence of CSR from loci with weak or absent population-genomic association highlights both the strength and limitation of this approach: machine learning can enrich for biologically relevant candidates, but it cannot fully capture context-dependent survival phenotypes without complementary functional validation. Finally, heterogeneity in clinical data sources and susceptibility testing conditions further bounds model performance and underscores the need for larger, better-annotated genomic datasets.

Beyond these technical considerations, our findings have broader conceptual implications for how antimicrobial resistance is defined and diagnosed. CSR occupies a space distinct from classical resistance, tolerance, or persistence as traditionally framed, emphasizing survival capacity during therapy rather than shifts in breakpoint-defined susceptibility or transient growth arrest. While MIC testing remains indispensable for guiding antibiotic choice, it captures only a narrow slice of bacterial response to antimicrobial stress. Our results demonstrate that genetically encoded, MIC-independent survival strategies can meaningfully influence treatment outcomes and should be considered when interpreting susceptibility data, designing predictive diagnostics, and developing therapeutic strategies.

In summary, this work reveals that *A. baumannii* can harbor clinically silent, genetically encoded survival adaptations that promote low level resistance during colistin therapy in vivo. By integrating machine learning with functional and in vivo validation, we identify CSR as a discrete survival layer that expands the conceptual boundaries of antimicrobial resistance. Recognizing and interrogating such hidden resistance states will be essential for improving diagnostic accuracy and mitigating treatment failure in the era of last-line antibiotics.

## Methods

### Pangenome Construction

AMR and genome data was downloaded from the BVBRC resource [37]. The pangenome was constructed using Roary (version 3.13.0) [38]. Input files consisted of annotated assemblies in GFF3 format produced by BAKTA [39]. The analysis was performed using default settings. The resulting output provided a gene presence/absence matrix and clusters of orthologous genes (COGs) for downstream analysis. Any COG with representative genes present in the ABUW strain were used for downstream analysis.

### Sequence Alignment of Gene Clusters

To investigate sequence variation within orthologous groups, amino acid sequences for identified gene clusters were extracted and aligned using ClustalW (version 2.0) [40]. Alignments were performed using the default parameters to ensure optimal residue matching across the gene clusters. These protein-level alignments were subsequently used for statistical comparisons to identify residue changes associated with antibiotic resistance.

### Statistical Analysis

Statistical associations were calculated using Python version 3.11 with the SciPy library (version 1.16). Fisher’s exact test was applied to the alignment data to test for significance. Contingency tables were constructed based on the presence or absence of specific amino acid residues between susceptible and resistant strains. P-values were calculated to determine statistical significance, with a threshold of p < 0.05.

### Bacterial strains and culture conditions

All strains were derived from *A. baumannii* AB5075-UW and obtained from the Manoil transposon mutant library [41]. Strains were cultured in cation-adjusted Mueller–Hinton broth (CAMHB; Becton Dickinson) at 37 °C with shaking at 200 rpm and stored as 30% glycerol stocks at −80 °C.

### Verification of transposon insertion sites

Insertion sites were confirmed by colony PCR using primers flanking each targeted locus [42]. Single colonies were lysed by heating at 95 °C for 10 min, and PCR was performed using Taq polymerase (New England Biolabs) under standard cycling conditions. Amplicons were resolved on 1% agarose gels and visualized with GelRed. Mutants producing bands approximately 900 bp larger than wild type were considered correctly disrupted. Clonality was verified by re-streaking and repeat PCR for strains with ambiguous results.

### Minimum inhibitory concentration (MIC) determination

Colistin MICs were determined by broth microdilution according to CLSI guidelines [43]. Overnight cultures were subcultured 1:100 in fresh CAMHB to an OD₆₀₀ of 0.4, diluted to OD₆₀₀ = 0.002, and inoculated into 96-well plates containing twofold serial dilutions of colistin sulfate. Plates were incubated at 37 °C for 18–20 h with shaking. MIC was defined as the lowest concentration showing no visible turbidity, and OD₆₀₀ readings were confirmed using a BioTek Synergy plate reader.

### Growth kinetics under colistin exposure

Overnight cultures were subcultured 1:100 into fresh CAMHB, grown to mid-log phase, and diluted into LB medium containing 0, 1, 2, or 4 µg ml⁻¹ colistin [44]. Growth was monitored at 37°C with continuous shaking, and OD₆₀₀ was recorded every 30 min for 24 h. Growth curves were normalized to antibiotic-free controls, and mutants were classified as wild type–like, intermediate, or clinically silent resistant based on relative growth dynamics.

### Membrane permeability assay

Membrane integrity was assessed by propidium iodide uptake [45]. Mid-log cultures were washed in PBS, adjusted to OD₆₀₀ = 0.2, incubated with 2 µM propidium iodide, and fluorescence (Ex 535 nm, Em 617 nm) was measured using a BioTek Synergy plate reader. Increased fluorescence relative to wild type indicated elevated permeability.

### Membrane potential assay

Membrane potential (Δψ) was measured using 3,3′-diethyloxacarbocyanine iodide [DiOC₂(3)] [46]. Cultures were adjusted to OD₆₀₀ = 0.2, stained with 10 µM dye, incubated 30 min in the dark, and fluorescence was measured at 488/525 nm. Lower fluorescence indicated depolarization; higher fluorescence indicated hyperpolarization.

### Efflux activity assay

Efflux activity was quantified by ethidium bromide accumulation [47]. Mid-log phase cells were washed twice in PBS, resuspended to OD₆₀₀ = 0.2 in PBS containing 2 µg ml⁻¹ ethidium bromide, and fluorescence (Ex 530 nm, Em 600 nm) was recorded every 60 s for 30 min. Lower fluorescence relative to wild type indicated increased efflux.

### Biofilm formation assay

Biofilm formation was quantified by crystal violet staining [48]. Overnight cultures were subcultured 1:100 into fresh CAMHB and incubated in 96-well plates for 24 h at 37 °C. Wells were washed with PBS, air-dried, stained with 0.1% crystal violet for 15 min, washed, and dye was solubilized in 95% ethanol. Absorbance at 590 nm was normalized to wild type.

### Oxidative-stress tolerance assay

Hydrogen peroxide sensitivity was assessed using a broth microdilution–like assay [49]. Twofold H₂O₂ dilutions in CAMHB were inoculated with cultures diluted to OD₆₀₀ = 0.002. Plates were incubated at 37 °C for 19 h, and growth inhibition relative to untreated controls was quantified.

### Murine pneumonia model

All animal experiments were approved by the University of Texas at Dallas IACUC. Female C57BL/6J mice (7–8 weeks old; n = 5/group) were anesthetized with ketamine/xylazine (10:1) and intratracheally inoculated with 1 × 10⁸ CFU of *A. baumannii* in 40 µl PBS [50,51]. Colistin was administered intraperitoneally at 0–20 mg kg⁻¹ two hours post-infection. After 24 h, lungs were harvested, homogenized, serially diluted, and plated for CFU enumeration. Bacterial burden was expressed as log₁₀ CFU mg⁻¹ tissue. Statistical significance was assessed by two-tailed paired t-test.

### Statistical analysis

All experiments were performed with at least three independent biological replicates unless otherwise stated. Data are reported as mean ± s.d. Statistical analyses were performed using GraphPad Prism (v10.0). Two-group comparisons used unpaired two-tailed t-tests; multiple comparisons used one-way ANOVA with Tukey’s post hoc test [52]. A significance cutoff of P < 0.05 was applied. In vivo CFU values were log₁₀-transformed prior to analysis.

## Acknowledgements

This work was supported in part by seed funding from The University of Texas at Dallas and by funding from the National institutes of Health (grants AI168159 and GM143053 to JMB). We thank the UT Dallas Animal Resource Center for assistance with animal husbandry and Dr. Xintong Dong for providing ketamine and xylazine.

---

**Extended Figure 1.**
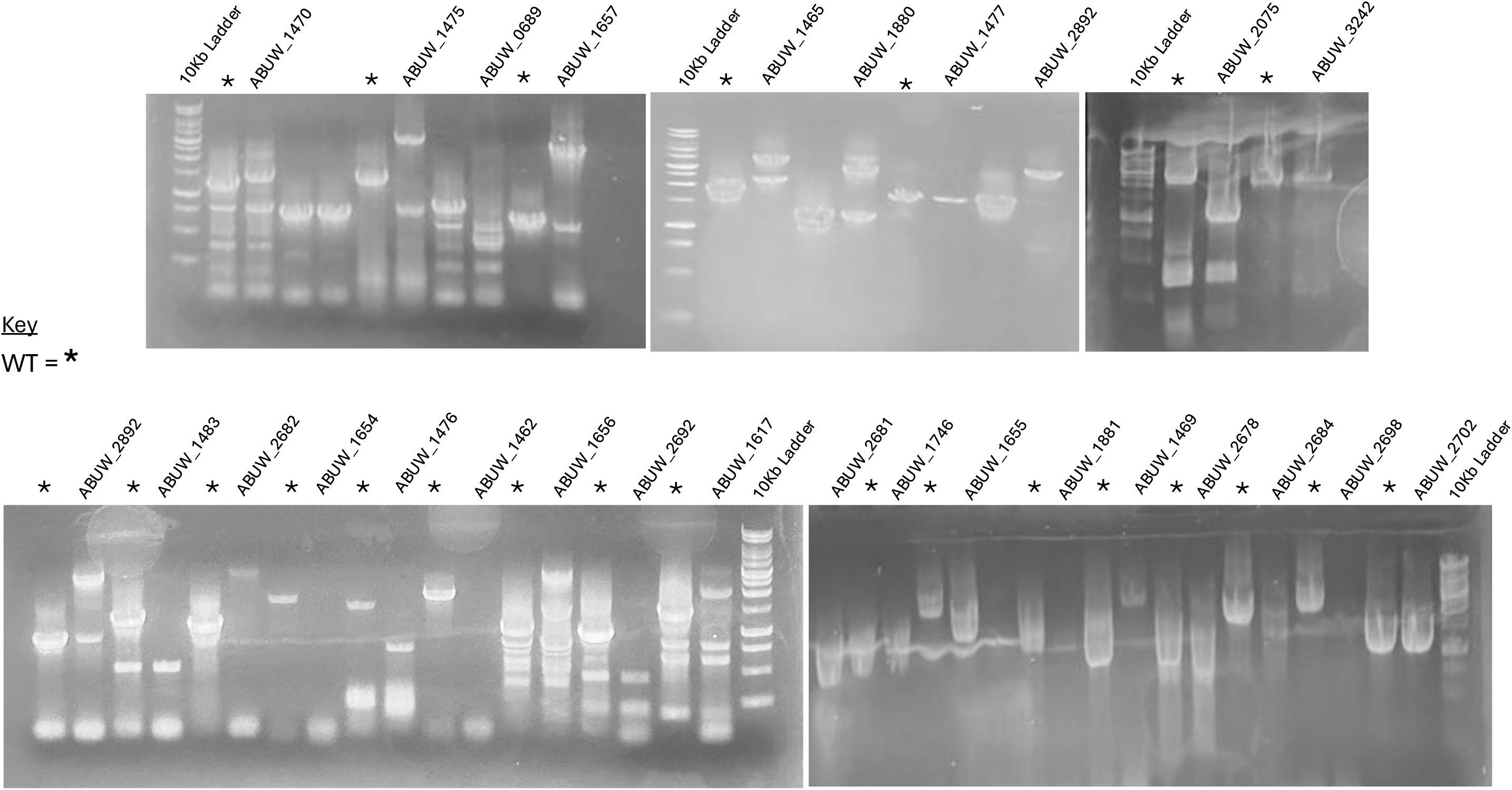
PCR validation of transposon insertion mutants. Representative agarose gel electrophoresis images showing PCR confirmation of transposon insertions in selected *A. baumannii* AB5075-UW mutants. Genomic DNA from individual transposon mutants was amplified using primer pairs flanking the predicted insertion sites and/or primers specific to the transposon and adjacent genomic regions. Correctly sized amplicons consistent with transposon insertion were observed for each mutant, while the wild-type control lacked the corresponding insertion-specific bands. Molecular weight markers (DNA ladders) are shown to indicate fragment sizes. These results confirm the presence and genomic location of transposon insertions in the validated mutant strains used for downstream phenotypic analyses.

**Extended Figure 2:**
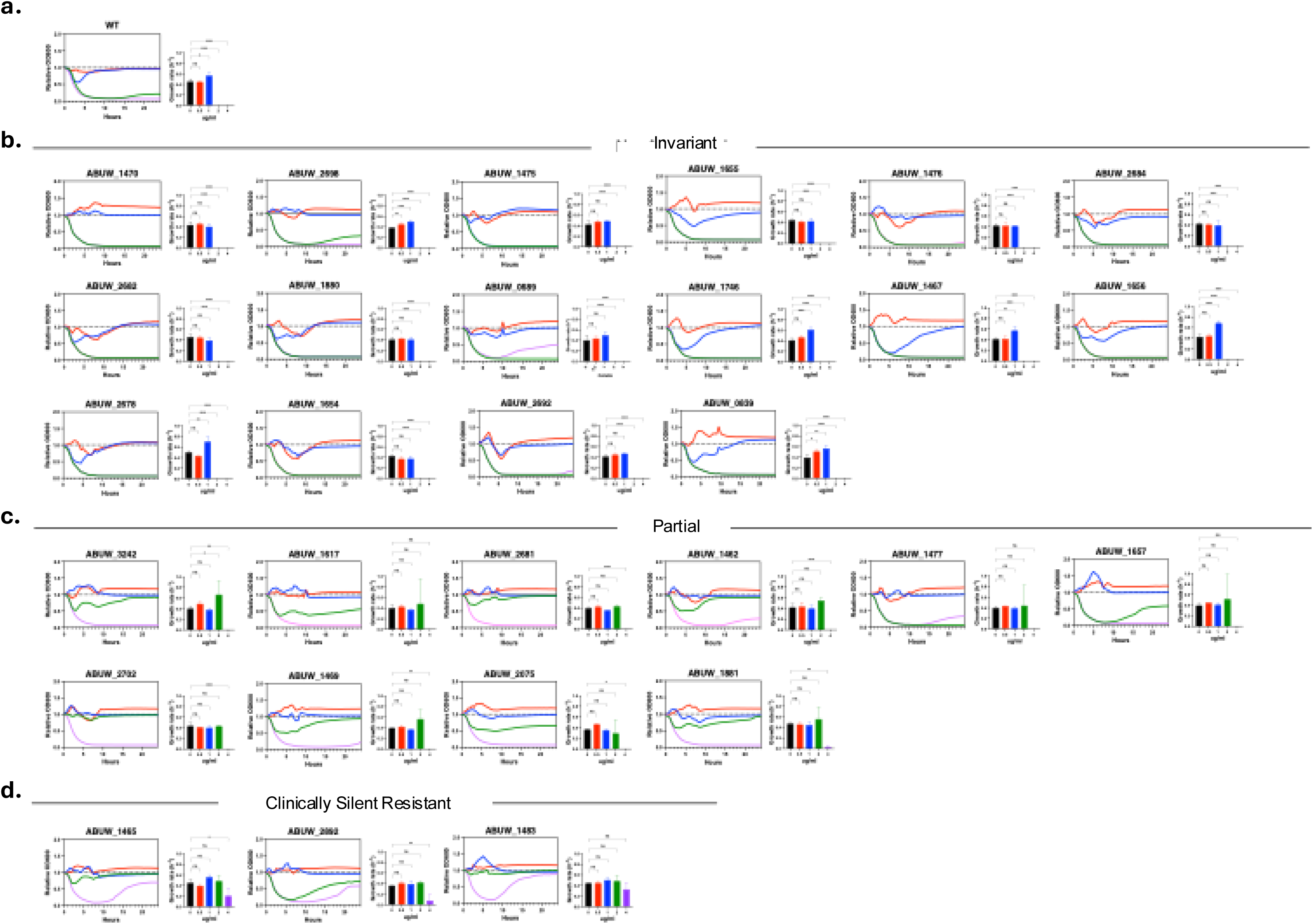
Complete growth-kinetics profiles of machine-learning–prioritized *A. baumannii* mutants under colistin exposure. Real-time growth kinetics were measured for all machine-learning–prioritized transposon mutants in the *A. baumannii* AB5075 background across increasing concentrations of colistin. Each panel displays optical density (OD₆₀₀) over time for an individual mutant cultured in the absence of colistin or under escalating colistin exposure, as indicated by color. Growth curves are shown relative to the wild-type strain assayed in parallel. Bar plots adjacent to each growth curve summarize endpoint growth or area-under-the-curve (AUC) metrics derived from the corresponding time-course data and are normalized to wild type. Mutants are ordered consistently with Figure 3. Panels are grouped by growth phenotype: (a) wild-type–like inhibition, (b) non-advantaged mutants, (c) intermediate growth recovery under colistin stress, and (d) clinically silent resistant mutants exhibiting sustained growth at concentrations inhibitory to wild type. This complete dataset provides the full growth-kinetic context underlying the representative phenotypes highlighted in the main figure.

**Extended Table 1:** Mutation-level association statistics for machine-learning–prioritized loci. This table summarizes mutation-level association metrics for genes prioritized by the supervised machine-learning framework. For each locus, the table reports the statistical significance of mutation enrichment ( *P* value), the number of resistant isolates harboring mutations at that locus, and the direction of association with colistin response.

| Gene Name | Gene Presence P-value | #Unique Mutations | Mean_%Resistant Strains Carrying Mutations | Mean_%_Susceptible Carrying Mutations | Odd Ratio |
| --- | --- | --- | --- | --- | --- |
| ABUW_1470 | 0.000000027 | 0 | 0 | 0 | 0 |
| ABUW_1475 | 0.0581 | 0 | 0 | 0 | 0 |
| ABUW_0689 | 0.0688 | 140 | 64.57142857 | 49.85983827 | 14.7115903 |
| ABUW_1657 | 0.0688 | 0 | 0 | 0 | 0 |
| ABUW_1465 | 0.0805 | 2 | 55 | 49.24528302 | 5.754716981 |
| ABUW_1880 | 0.0805 | 0 | 0 | 0 | 0 |
| ABUW_1477 | 0.0805 | 0 | 0 | 0 | 0 |
| ABUW_2892 | 0.153 | 0 | 0 | 0 | 0 |
| ABUW_1483 | 0.153 | 0 | 0 | 0 | 0 |
| ABUW_2682 | 0.17 | 0 | 0 | 0 | 0 |
| ABUW_1654 | 0.289 | 6 | 35 | 21.19496855 | 13.80503145 |
| ABUW_1476 | 0.289 | 0 | 0 | 0 | 0 |
| ABUW_1462 | 0.289 | 0 | 0 | 0 | 0 |
| ABUW_1656 | 0.293 | 4 | 50 | 49.90566038 | 0.094339623 |
| ABUW_2692 | 0.332 | 0 | 0 | 0 | 0 |
| ABUW_1617 | 0.344 | 356 | 64.74719101 | 49.70850117 | 15.03868985 |
| ABUW_2681 | 0.408 | 0 | 0 | 0 | 0 |
| ABUW_1746 | 0.408 | 0 | 0 | 0 | 0 |
| ABUW_1655 | 0.44 | 5 | 30 | 20.75471698 | 9.245283019 |
| ABUW_1881 | 0.44 | 0 | 0 | 0 | 0 |
| ABUW_1469 | 0.473 | 0 | 0 | 0 | 0 |
| ABUW_2678 | 0.473 | 0 | 0 | 0 | 0 |
| ABUW_2684 | 0.505 | 309 | 15.14563107 | 7.279721561 | 7.865909507 |
| ABUW_2698 | 0.505 | 1 | 0 | 28.30188679 | -28.30188679 |
| ABUW_2702 | 0.666 | 0 | 0 | 0 | 0 |
| ABUW_1467 | 0.697 | 2 | 30 | 6.603773585 | 23.39622642 |
| ABUW_0839 | 0.85 | 68 | 63.67647059 | 48.54051054 | 15.13596004 |
| ABUW_2075 | 0.923 | 159 | 59.81132075 | 46.34626795 | 13.46505281 |
| ABUW_3242 |  | 0 | 0 | 0 | 0 |

**Supplementary Table 1:** Global gene presence/absence analysis (xlxs sheet)

